# Autoprot: Processing, Analysis and Visualization of Proteomics Data in Python

**DOI:** 10.1101/2024.01.18.571429

**Authors:** Julian Bender, Wignand W. D. Mühlhäuser, Johannes P. Zimmerman, Friedel Drepper, Bettina Warscheid

## Abstract

The increasing numbers of complex quantitative mass spectrometry-based proteomics data sets demand a standardised and reliable analysis pipeline. For this purpose, Python-based analysis, particularly through Jupyter notebooks, serves as a simple yet powerful tool. Nevertheless, the availability of Python software for standardised and accessible MS data analysis is limited, and this software is often constrained to using analysis functions written in Python. This excludes existing and well-tested software, for example written in R. Despite this, Python offers several interactive data visualisation modules that greatly enhance exploratory research and facilitate result communication with collaboration partners. Consequently, there is a need for an integrated and Jupyter-compatible Python analysis pipeline that incorporates R algorithms and interactive visualization for proteomics data analysis.

**Summary:** We developed autoprot, a Python module for simplified analysis of quantitative mass spectrometry-based proteomics experiments processed with the MaxQuant software. It provides access to established functions written in both Python and R for statistical testing and data transformation. Moreover, it generates JavaScript-based interactive plots that can be integrated into interactive web applications. Thereby, autoprot offers standardised, fast and reliable proteomics data analysis while maintaining the high customisability required to tailor the analysis pipeline to specific experiments.

**Availability and Implementation:** Autoprot is implemented in Python ≥ 3.9 and can be downloaded from https://github.com/ag-warscheid/autoprot. Online documentation is available at https://ag-warscheid.github.io/autoprot/.

## 1. Introduction

Advances in high-resolution protein mass spectrometry (MS) and the increasing availability of this high-end technology for labs addressing a large variety of biomolecular and biomedical questions have expanded the number, size, and complexity of the acquired MS raw files for subsequent data analysis. Accordingly, various software platforms have been developed to analyse and quantify peptides and proteins from MS raw files generated from data-dependent or data-independent tandem MS (MS/MS) measurements. Here, the freely available software MaxQuant (Cox and Mann 2008; Cox et al. 2009, 2011; Tyanova, Temu and Cox 2016) in particular has matured to a reference pipeline for data-dependent proteomics studies. Furthermore, to analyse proteomics data from data-independent acquisition (DIA) the software MaxDIA has been recently implemented in MaxQuant (Sinitcyn et al. 2021). However, programs for the analysis of MaxQuant output tables generally exist between two extremes: Simple, intuitive tools of limited functional scope, like Microsoft Excel, and advanced approaches requiring a certain proficiency of the user in a programming language. Tools trying to bridge that gap include Perseus (Tyanova et al. 2016), a graphical user interface for statistical analysis of MaxQuant data, statistical analysis in the cloud-based galaxy-framework (Pinter et al. 2022), and various code packages for streamlining the analysis workflow (Heming et al. 2022: 2; Sueur et al. 2023).

In our group, we found that Python code generated in Jupyter notebooks is well suited for both an advanced programmer and a beginner in proteomics data analysis. It provides the necessary flexibility to analyse tabular files resulting from a variety of proteomics workflows and biological questions. Moreover, large proteomics data sets often require unbiased explorative analysis. To address this, we use (i) the plotly package, which integrates well with Jupyter notebooks and produces meaningful interactive plots and figures as well as (ii) plotly-based dashboards (Hossain 2019), which make large proteomics data sets easily accessible and comprehensible for collaboration partners. Additionally, a centralised Jupyter server enables to maintain a complete programming environment by a single skilled user, thereby avoiding the need of installing software by the general user. However, despite Python being beginner friendly and generally well adopted, specialised algorithms for the statistical analysis of omics datasets are often exclusively available in the R programming language. Based on these considerations, we developed autoprot, a Python software package that integrates well into Jupyter notebooks, generates interactive figures and dashboards, and utilises advanced statistical functionality from R.

## 2. Implementation

Autoprot serves as an interface between the user and various data processing analysis algorithms (see **Table S1** for details). It utilises Python syntax and relies on well-established data analysis packages like pandas and NumPy. Advanced statistical R algorithms are invoked through a dedicated R installation interfaced by Python’s subprocess routine. This ensures that autoprot functions reliably under all operating systems on which R is running and provides Python users an easy way to employ powerful statistical tools otherwise exclusive for R. Autoprot is designed to focus on the three main parts of proteomics MS data analysis: pre-processing, analysis, and visualisation (**Figure 1**).

**Figure 1:**
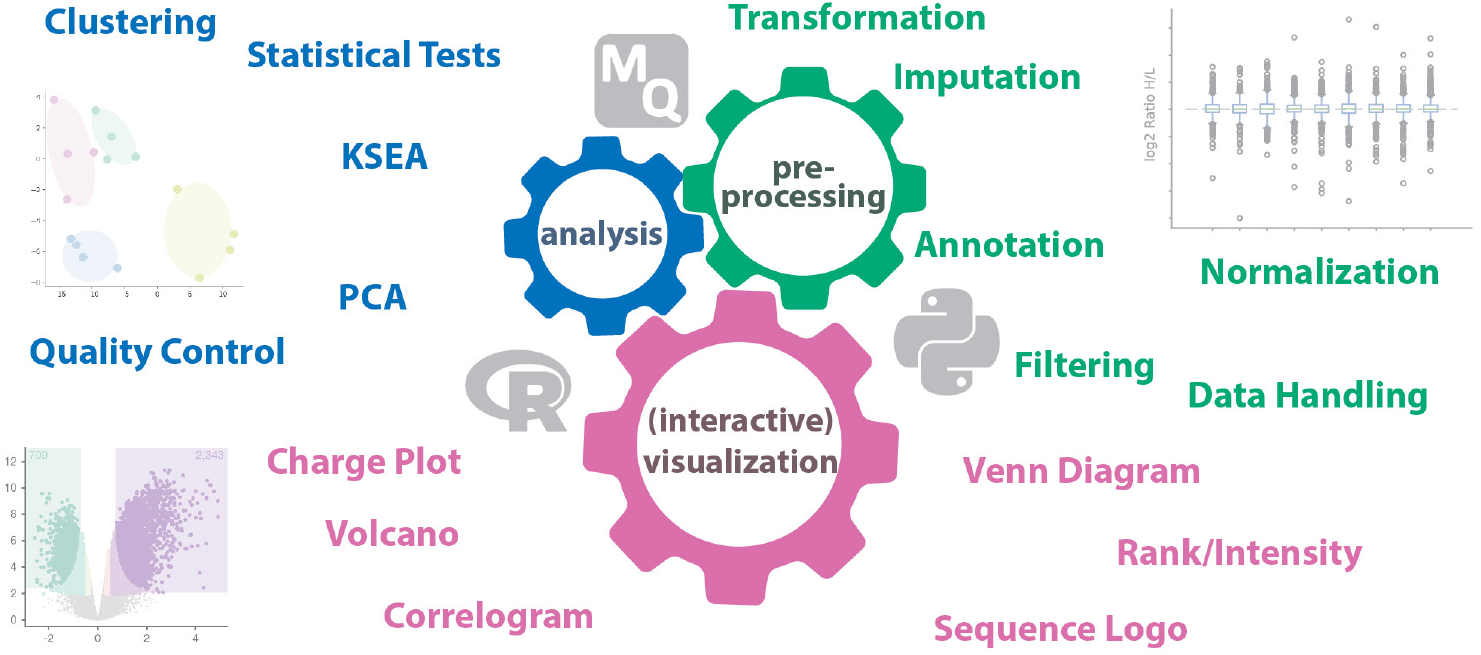
Autoprot functionalities. The package combines Python and R functions for the analysis of MaxQuant (MQ) search results. It is divided into three main modules: analysis (blue), preprocessing (green) and interactive visualization (pink) containing various functions with the indicated objectives (color-coded). KSEA: Kinase-substrate enrichment analysis (Wiredja et al. 2017); PCA: Principal component analysis.

### 2.1. Pre-processing

The output tables from MaxQuant are generally not suited in their original format for data analysis but require some curation and transformation. For example, MaxQuant provides raw intensities which must be log-transformed for statistical analysis. MaxQuant also generates semicolon-separated fields containing multiple protein identifiers that can be unstacked in autoprot. Some data sets require filtering by sequence coverage, phosphosite localisation probability, valid values in multiple replicates or GO annotation (Gene Ontology Consortium et al. 2023). Common data analysis techniques such as clustering require that the input is normalised and missing values are removed. Here, autoprot offers a set of functions (including quantile normalisation, variance-stabilising normalisation, and cyclic LOESS) all sharing the same call structure so that they are easily interchanged in an analysis script. As implementation for variance-stabilising normalisation, the R package “vsn” from Bioconductor (Huber et al. 2002) was chosen as it is tuned to omics data and is widely accepted in the field. For data imputation, i.e., the replacement of values missing either at random or through insufficient peptide intensity, autoprot offers access to advanced methods such as data-dependent selection of an imputation algorithm (Egert et al. 2021).

### 2.2. Analysis

The control of proteomics MS data quality becomes increasingly important with data size and complexity. Therefore, the analysis submodule contains quality control functions to assess missed cleavages, enrichment of post-translationally or chemically modified peptides, metabolic or chemical stable isotope labelling efficiency or missing values between replicates. The submodule contains functions for comparative statistical analysis such as the t-test but also incorporates more advanced approaches including linear models and rank sum tests. Other advanced methods, which usually require a multi-step approach, are organized in dedicated classes. For example, principal component analysis is included in the AutoPCA class that streamlines data analysis and validation of the results through visual inspection. Similarly, we included hierarchical clustering and K-means clustering as specific classes into the analysis submodule. The clustering classes implement complete workflows that, for example, consist of hierarchical clustering, the generation of linkages, determining the optimal number of clusters and the final clustering. These steps are separately accessible to allow stepwise fine-tuning of the data analysis to obtain most informative results. In general, autoprot analysis classes simplify tasks that would otherwise require numerous lines of code. Similar functions in autoprot follow a common argument structure, thereby facilitating the exchange and comparison of e.g., one clustering method against another. For functional interpretation of certain protein subsets, the analysis module contains functions for Gene Ontology (GO) annotation and kinase-substrate enrichment analysis (KSEA; Wiredja, Koyutürk and Chance 2017). The latter is implemented as a class providing a streamlined workflow from annotation to visualisation (Reimann et al. 2020; Fricke et al. 2023)

### 2.3. Visualisation

Drawing functional conclusions from quantitative MS data is aided by reliable and conclusive visualisation. We integrated a set of commonly employed visualisations for large proteomics data into the visualisation module of autoprot. These include volcano plots, Venn diagrams, boxplots, sequence logos, or intensity rank plots. All visualisation functions come with an interactive version based on the plotly library (the interactive versions have the same name as the functions that generate static plots with an additional letter “i” as prefix, e.g., volcano and ivolcano). Thus, users can easily switch between static plots (e.g., for presentations) and interactive data exploration. Moreover, the figure object returned by the interactive versions can be incorporated into interactive web dashboards thereby simplifying the path from data analysis to data sharing.

## 3. Examples

In this study, we illustrate the application of autoprot by creating four Jupyter notebooks (refer to the Data Availability section for the link to the GitHub repository). These notebooks utilize autoprot to reanalyse publicly accessible proteomics data obtained from the PRIDE repository (Perez-Riverol et al. 2022). The notebooks cover various analyses, including: (1) utilizing autoprot for the examination of affinity purification MS results through diverse statistical methods and interactive data visualization for inspection; (2) employing autoprot for the analysis of stable isotope-labelling in cell culture (SILAC) experiments, encompassing channel normalization, principal component analysis, and statistical analysis; and (3) showcasing the capability of autoprot for the analysis of phosphoproteomics data, focusing on working with peptide-level information, conducting missing value analysis, performing functional annotation via KSEA, and implementing clustering. Each workflow is comprehensively documented within the respective notebooks.

For brevity, we focus here on the analysis of a dataset created before autoprot (or its predecessors) were developed: a dimethyl-labelling based analysis of the proteome of C2C12 skeletal myocytes before and after differentiation of myoblasts to myotubes and with and without mild electrical pulse stimulation (EPS) for the in vitro formation of contracting cross-striated myofibrils (Reimann et al. 2017). When analysing the ratios between myoblasts and myotubes using a conventional t-test **(Figure 2**, top), no significantly regulated proteins are identified based on the MaxQuant ratios. The same is true for the protein ratios between EPS-treated myotubes and myoblasts and between EPS-treated myotubes and myotubes without EPS. Analysing the same ratios with linear modelling and considering the different isotope labels, the resulting p-values are much lower (**Figure 2**, middle). Moreover, mapping the clusters of differentially regulated proteins from the publication on the volcano plots, proteins from a cluster attributed to sarcomeric proteins produced during differentiation (red) and nuclear proteins that are less expressed in differentiated cells (blue) shows a clear correlation of the original annotation with the statistical analysis in autoprot. Lastly, we also reproduced the whole clustering analysis in autoprot (**Figure 2**, bottom) showing a similar result as previously published. Of note, the plots shown in **Figure 2** are shown as returned from autoprot without any post-processing.

**Figure 2.**
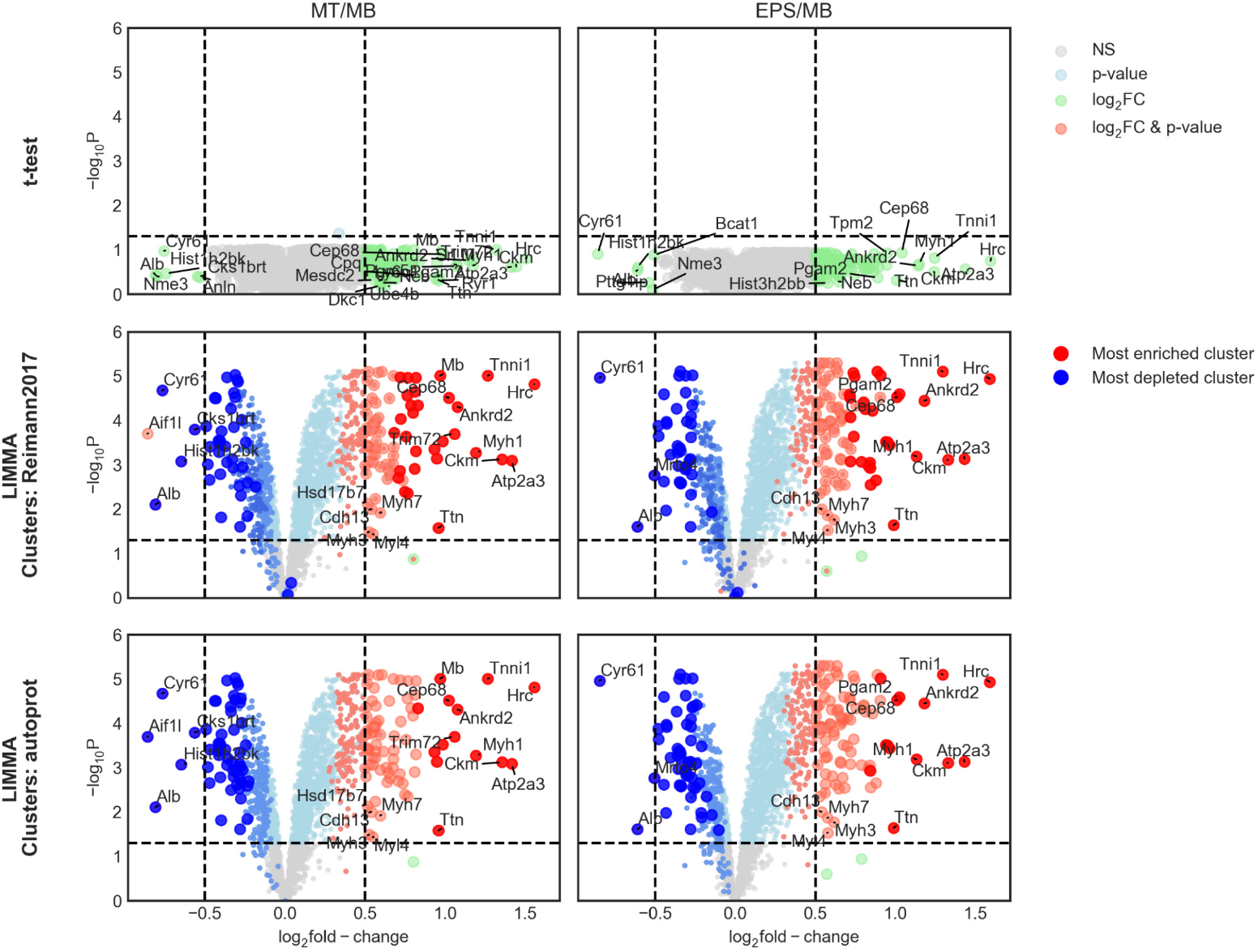
Autoprot analysis of the myofibrillar proteome. Ratios from chemically labelled myoblast (MB) and myotube (MT) proteins before and after mild electrical pulse stimulation (EPS) from Reimann et al. (2017) were analyzed with autoprot. Conventional t-test statistics (top) does not produce significantly regulated genes during differentiation. Linear modelling available through autoprot (middle) detects significant changes in the proteome and significance level correlates with clusters 1 and 2 (red) and clusters 9-11 (blue) from the original publication. Cluster analysis in autoprot (bottom) recapitulates the cluster annotation.

## 4. Conclusion

With autoprot, we have created a tool for standardised access to all elements in the proteomics MS data analysis chain, spanning from pre-processing to visualisation. By integrating analysis functions from Python and R and making the results accessible in an interactive manner, autoprot significantly accelerates the analysis of various kinds of proteomic datasets (e.g., phosphoproteomics, LFQ, SILAC, etc). Leveraging Jupyter notebooks and a centralized Jupyter Hub infrastructure eliminates the need for users to install and update software, thereby enhancing ease of use. Furthermore, it reduces the entry barrier for using Python as a robust routine tool for advanced proteomics data analysis, making it accessible even to the inexperienced user. Therefore, we consider autoprot to be an extremely valuable and broadly applicable tool for analysing complex and large quantitative proteomics MS data, enabling the generation and provision of meaningful and dependable proteomics results of high quality.

## 6. Data availability

Autoprot is published as open-source package and the source code together with example notebooks showcasing the use of autoprot is publicly available on GitHub (https://github.com/ag-warscheid/autoprot). The code for the example shown here is available at examples/04_ratio_clustering.ipynb

## Acknowledgements

We thank Hirak Das for extensive testing and Silke Oeljeklaus for constant feedback on the usefulness of the functions. BioRxiv Word template from https://github.com/chrelli/bioRxiv-word-template.

## SUPPLEMENTARY DATA

**Table S1**: Autoprot functions

